# Identification of an anti-CRISPR protein that inhibits the CRISPR-Cas type I-B system in *Clostridioides difficile*

**DOI:** 10.1101/2023.05.22.541795

**Authors:** Polina Muzyukina, Anton Shkaruta, Noemi M. Guzman, Jessica Andreani, Adair L. Borges, Joseph Bondy-Denomy, Anna Maikova, Ekaterina Semenova, Konstantin Severinov, Olga Soutourina

**Author notes:** University of Melbourne, Parkville, Victoria 3010, Australia (Anton Shkaruta). Synthetic Biology, Institut Pasteur, Université Paris Cité, CNRS UMR 6047, 75015 Paris, France.

## Abstract

CRISPR-Cas systems provide their prokaryotic hosts with adaptive immunity against mobile genetic elements. Many bacteriophages encode anti-CRISPR (Acr) proteins that inhibit host defense. The identification of Acr proteins is challenging due to their small size and high sequence diversity, and only a limited number has been characterized to date. In this study, we report the discovery of a novel Acr protein, AcrIB2, encoded by the φCD38-2 *Clostridioides difficile* phage that efficiently inhibits interference by the type I-B CRISPR-Cas system of the host and likely acts as a DNA mimic. Most *C. difficile* strains contain two *cas* operons, one encoding a full set of interference and adaptation proteins and another encoding interference proteins only. Unexpectedly, we show that only the partial operon is required for interference and is subject to inhibition by AcrIB2.

## Introduction

Competition for survival in nature drives organisms to continuously adapt and evolve, leading to the evolution of species over time (1, 2). A constant battle between prokaryotes and parasitic mobile genetic elements (MGEs), most notably viruses, provides a vivid illustration of this principle. To avoid extermination by viruses, which are estimated to outnumber their prokaryotic hosts by an order of magnitude (3), cells have evolved numerous defense strategies. In their turn, to avoid extinction, phages have evolved countermeasures to overcome specific defenses of their hosts. Adaptive prokaryotic CRISPR-Cas immunity systems have received much attention due to their unique mechanism of action and significance for biotechnology and biomedicine. These RNA-guided defenses consist of CRISPR arrays and associated *cas* genes. During CRISPR adaptation, the hosts integrate short sequences derived from infectious agents’ genomes as spacers into the CRISPR arrays. During CRISPR interference, the Cas proteins guided by short CRISPR RNAs (crRNAs) transcribed from the array recognize and eliminate invading pathogens genomes with sequences complementary to crRNA spacers (4–6).

One way in which bacteriophages and other MGEs can evade CRISPR-Cas immunity is by modifying or removing targeted DNA sequences from their genome (7–9). However, this strategy has limitations, particularly when CRISPR-Cas targets essential regions of MGE genomes. Another strategy is to avoid recognition by CRISPR-Cas (and other DNA-targeting host defenses, such as restriction-modification systems) by extensively modifying the invader’s DNA or creating excluded compartments in infected cells that make invader DNA inaccessible to host defense systems (9–11). Yet another common strategy relies on anti-CRISPR proteins (Acrs) that are encoded by MGEs and inhibit CRISPR-Cas immunity by diverse mechanisms (12).

The number of identified and experimentally characterized Acrs is steadily growing (13) and is constantly updated (tinyurl.com/anti-crispr). Known Acrs inhibit CRISPR interference by preventing target binding, target cleavage, or crRNA interaction with Cas interference proteins (14). Most Acrs are small proteins, and many have highly negative overall charge, likely acting as DNA mimics (15–18).

Within the phage genomes, known *acr* genes are often paired with anti-CRISPR-associated (*aca*) genes. The Aca proteins are helix-turn-helix (HTH) domain-containing transcription factors that regulate *acr* transcription (19). Genes coding for small proteins with the AP-2 DNA-binding domain are also frequently observed in *acr* loci (20). While the diversity of Acrs poses a significant challenge for their identification by means of bioinformatics (21), the “guilt-by-association” approach involving analysis of sequences flanking *aca*-like genes has met with considerable success (19, 22). Another strategy involves the analysis of prokaryotic genomes containing CRISPR arrays with spacers matching sequences in a host’s own genome. In these cases, self-immunity is often prevented by Acrs encoded in prophages (19).

Interest in the discovery of new Acr proteins is driven by a potentially wide range of applications that they could serve, including the development of phage therapy for pathogenic bacteria (14). A phage that can efficiently overcome host CRISPR-Cas defense by employing Acrs would clearly be a preferred candidate for therapeutic application. Phage therapy is considered a promising alternative to antimicrobial treatments against the widespread anaerobic spore-forming bacterium *Clostridioides difficile*, which poses a significant threat to human health all over the world (23–25). The type I-B CRISPR-Cas system of *C. difficile* is highly active and limits infection by phages (23, 26–28). Apart from *in silico* predictions, no anti-CRISPR proteins that target type I-B CRISPR-Cas systems have been characterized yet (29).

In this paper, we report a discovery of a new Acr protein that inhibits interference by the *C. difficile* CRISPR-Cas. This protein, which we name AcrIB2, is encoded by a temperate *C. difficile* phage φCDHM38-2. Sequence analysis indicates that proteins similar to AcrIB2 may be common to other clostridial phages. Most *C. difficile* strains encode two sets of type I-B *cas* genes. We show that the products of one *cas* gene set play no role in CRISPR interference, at least in laboratory settings. Consequently, it follows that AcrIB2 targets CRISPR interference encoded by proteins encoded by the remaining active *cas* genes set. Counterintuitively, the functional set of *cas* genes is incomplete, i.e., lacks genes required for CRISPR adaptation, while the apparently non-functional set has the full complement of interference and adaptation genes. These findings thus may hint at functional specialization between the duplicated *cas* operons of *C. difficile,* the nature of which remains to be determined.

## Results

### Search for putative anti-CRISPR loci in the genomes of *C. difficile* bacteriophages

Previously, while searching for homologs of AcrIC5, a phage inhibitor of type I-C CRISPR-Cas system from *Pseudomonas delhiensis*, León *et al.* discovered a 66 amino acid long hypothetical *Cryptobacterium curtum* protein of an unknown function (30). This protein was 63% identical to AcrIC5. The gene coding for this protein is adjacent to a gene encoding a 196 amino acid protein with a predicted AP2 DNA-binding domain. Genes encoding AP2 domain proteins are frequently observed in *acr* loci (20). We found a *Clostridium sp.* gene encoding a 159 amino acid AP2 domain protein that shared 30% sequence identity with *Cryptobacterium curtum AP2* domain protein (Fig. 1A). Using the *Clostridium sp.* sequence as a query, genes encoding highly similar AP2 domain proteins were found in the genome of *C. difficile LIBA2811* and in *C. difficile* phage φCDHM13 (Fig. 1A). The φCDHM13 gene is annotated as *gp27* and has no assigned function (31). Immediately upstream of the *gp27*, *gp26*, also a gene of unknown function, is located. A corresponding gene is also found in *C. difficile LIBA2811*. *gp28* and *gp29*, genes located immediately downstream of φCDHM13 *gp27*, partially overlap. Their homologs in *C. difficile LIBA2811* are fused. We hypothesized that the products of *gp26* and/or *gp28/gp29* might function as anti-CRISPR proteins targeting the *C. difficile* I-B CRISPR-Cas system. The *gp27* may function as an Aca protein.

**Figure 1.**
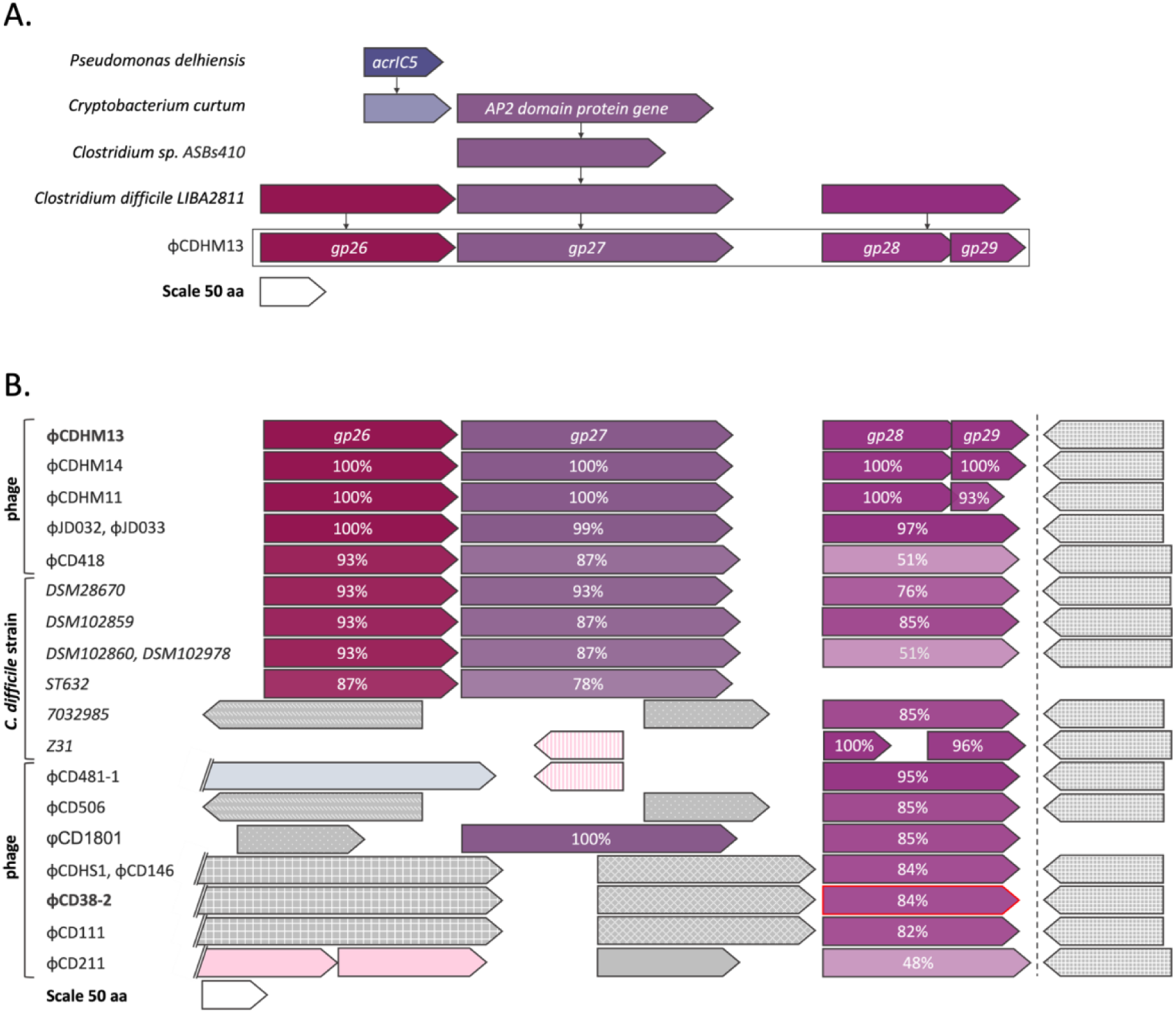
Putative anti-CRISPR loci of clostridial phages. Genes are represented by arrowed boxes drawn to scale (a scale is shown at the bottom of each panel). (A) A gene coding for an AP2 domain protein is located downstream of a homolog of *Pseudomonas delhiensis acrIC5* in *Cryptobacterium curtum*. A homolog of *C. curtum* AP2 domain protein coding gene was found in *Clostridium sp*., leading to the identification of a potential anti-CRISPR locus centered around the *AP2* domain protein gene in *C. difficile* strain *LIBA2811* and phage φCDHM13. (B) Using phage φCDHM13 *gp27* gene as a query, corresponding sequences from other clostridial phages and prophages were retrieved. Homologous genes are shown by matching colors and percentage of identity to corresponding φCDHM13 gene products is indicated. Genes whose products are non-homologous to φCDHM13 are colored grey. Genes denoted by a pink color encode potential transcriptional regulators or proteins containing HTH domain. Grey- and pink-colored genes sharing high sequence similarity are indicated with the same pattern. Light blue colored gene encodes amidase, a protein associated with a lysis module. The names of the two phages whose putative anti-CRISPR proteins were tested for function are highlighted in bold font. The φCD38-2 gene identified as *acrIB2* gene in this work is indicated by a red outline.

Subsequent bioinformatic analysis revealed that homologs of putative anti-CRISPR proteins are encoded by some other clostridial prophages and phages. Similarly to the case of *C. difficile LIBA2811*, some phages encoded φCDHM13 *gp28-gp29* fusions (Fig. 1B). Genes coding for such fused proteins were mostly found in phages that did not encode homologs of φCDHM13 *gp26* and *gp27* (for example, φCD38-2). Instead, they contained short open reading frames coding for proteins of unknown function (grey in Fig. 1B).

### Experimental validation of AcrIB2 from phage φCD38-2 as an inhibitor of *C. difficile* CRISPR-Cas interference

To assess the activity of predicted Acr proteins, each of the four genes from the predicted *acr* locus of phage φCDHM13 and the fusion of φCDHM13 gp28 and gp29 homologs from phage φCD38-2 was cloned, under the control of inducible pTet promoter, in a derivative of conjugative plasmid pRPF185Δ*gus* (26, 32). The only difference of the cloning vector from pRPF185Δ*gus* was the presence of a protospacer matching the first spacer of the *C. difficile* 630Δ*erm* CRISPR3 array. The cloned protospacer also contained a consensus CCA PAM sequence. Previous experiments showed that the *C. difficile* 630Δ*erm* CRISPR-Cas prevents conjugation of pRPF185Δ*gus* derivative carrying the protospacer (28). We, therefore, reasoned that a plasmid-borne inhibitor of CRISPR interference might restore conjugation efficiency. The original pRPF185Δ*gus* and its derivative carrying the protospacer only were used as controls. Transconjugants were selected on plates supplemented with thiamphenicol (Tm, pRPF185Δ*gus* provides cells with resistance to this antibiotic) and ATc to induce the expression of cloned phage genes. In agreement with published data (28), no transconjugants were observed with protospacer-containing pRPF185Δ*gus* plasmid. None of the φCDHM13 genes tested, including the *gp28-gp29* pair encoding the putative split Acr, restored conjugation efficiency (data not shown). However, conjugation of a plasmid expressing the fused homolog of φCDHM13 *gp28-29* from φCD38-2 was partially restored (Supplementary Fig. 1). We, therefore, concluded that the φCD38-2 protein acts as an anti-CRISPR and named it AcrIB2.

The partial effect of AcrIB2 on conjugation efficiency may be due to the fact that CRISPR interference with pre-existing crRNA produced from the first spacer of CRISPR3 array in the recipient cell may occur before the synthesis of sufficient amounts of AcrIB2 takes place. To overcome this limitation, we designed a different strategy that relies on a plasmid carrying an ATc-inducible mini CRISPR array containing a spacer that targets the *C. difficile hfq* gene (Fig. 2A). Elsewhere, we show that induction of mini CRISPR array transcription leads to cleavage of genomic DNA by the endogenous CRISPR-Cas system of *C. difficile*, therefore, preventing conjugation (33). We reasoned that if the self-targeting plasmid contains an ATc-inducible *acrIB2* gene, the anti-CRISPR activity of AcrIB2 will inhibit self-cleavage, leading to the appearance of transconjugants (Fig. 2A). Accordingly, *C. difficile* 630Δ*erm* transconjugants harboring various plasmids were obtained in the absence of induction, and their growth in liquid cultures in the presence or in the absence of the ATc inducer was monitored. As can be seen from Fig. 2B, the growth of induced culture harboring the self-targeting plasmid was strongly inhibited. In contrast, cells harboring a self-targeting plasmid and the *acrIB2* gene grew as fast as control cells carrying the empty vector. The number of CFUs in the cultures was determined at various times post-induction. As can be seen from Fig. 2C, as early as 1 hour post-induction, the number of viable cells in the culture harboring the self-targeting plasmid dropped 4 orders of magnitude compared to the uninduced control.

**Figure 2.**
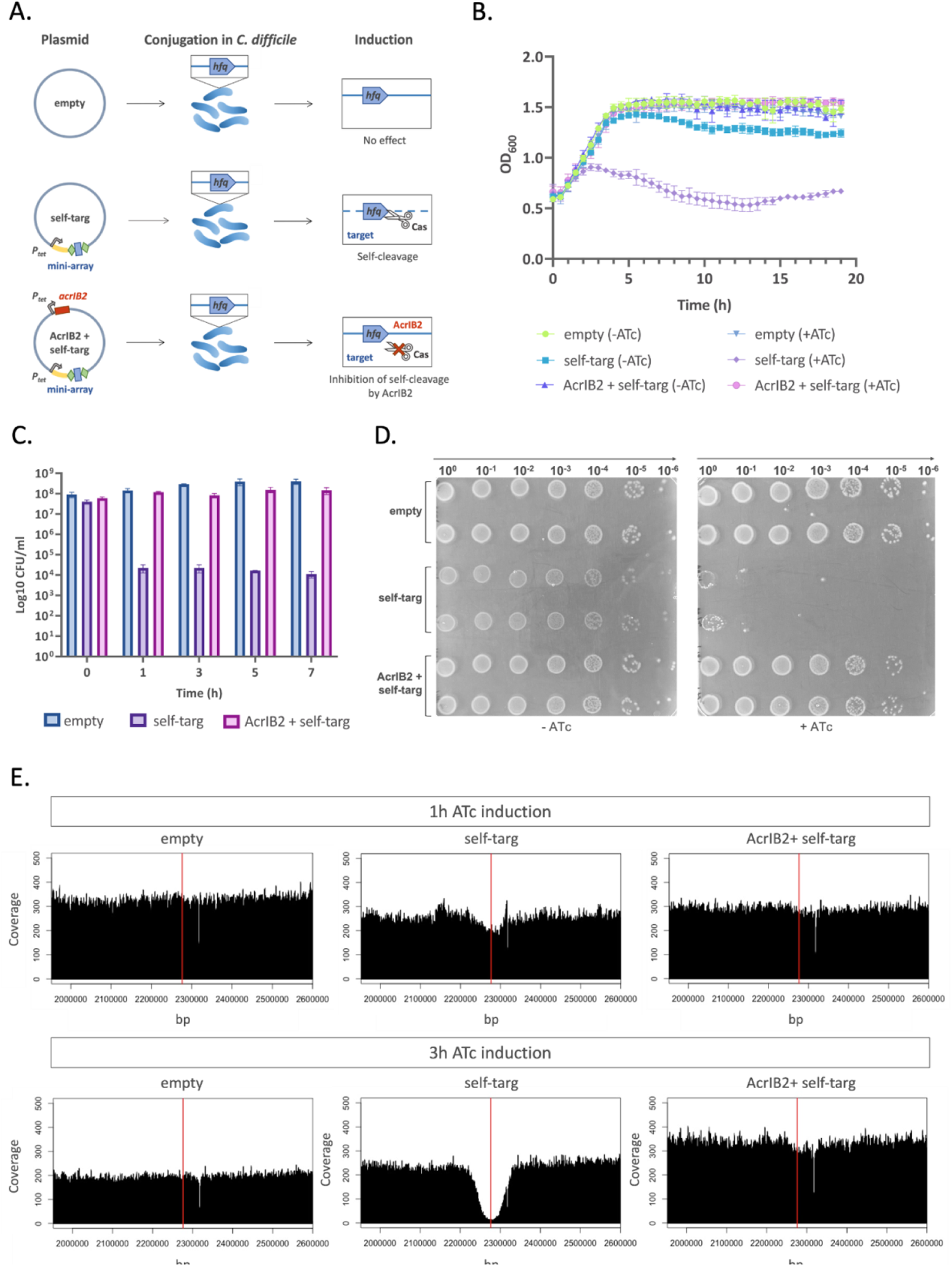
Anti-CRISPR protein AcrIB2 inhibits CRISPR interference in *C. difficile.* (A) A self-targeting strategy to reveal anti-CRISPR activity of plasmid-borne genes relies on a plasmid that carries, under the control of an inducible Ptet promoter, a mini CRISPR array with a spacer that targets the *C. difficile hfq* gene. Green rhombi indicate CRISPR repeats, the blue rectangle indicates a spacer, the leader sequence is indicated in yellow. “Self-targ” stands for self-targeting plasmid, “AcrIB2 + self-targ” stands for self-targeting plasmid with an ATc-inducible *acrIB2* gene. The control plasmid is referred to as “empty”. (B) The effect of anti-CRISPR on self-targeting inhibition of bacterial growth in liquid BHI medium supplemented with thiamphenicol (Tm, selects for cells carrying plasmids) in the presence or in the absence of the ATc inducer. Plotted values represent means, and error bars represent the standard error of the means (N = 3 biologically independent samples). (C) *C. difficile* cultures were grown in liquid BHI medium supplemented with Tm and induced with ATc. At indicated times post-induction, log_10_CFU/mL was determined by plating serial dilutions of cultures on BHI agar with Tm only. Values represent means, and error bars represent the standard error of the means (N = 3 biologically independent samples). (D) Aliquots of tenfold serial dilutions of *C. difficile* cultures conjugated with indicated plasmids were deposited on the surface of BHI agar supplemented with Tm with or without the ATc inducer. A representative result from at least three independent experiments is shown. (F) The effect of self-targeting/its inhibition by AcrIB2 on genomic DNA content as revealed by coverage of a fragment of *C. difficile* genome containing the self-targeted protspacer with ILLUMINA sequencing reads. The red vertical line indicates the location of the protospacer.

The same result was obtained when serial dilutions of aliquots of uninduced transconjugant cultures were spotted on plates with or without ATc. As can be seen from Fig. 2D, colony formation by cells harboring the self-targeting plasmid in the presence of ATc was severely impaired. In contrast, cultures harboring the self-targeting plasmid with *acrIB2* or the empty vector plasmid contained the same number of viable cells both in the presence and in the absence of the inducer. While rare colonies that formed in the places where drops of concentrated cultures of cells harboring the self-targeting plasmid were not studied systematically, we assume that they are escapers that contain mutations in the CRISPR-Cas system of the host, the targeted protospacer of the host, or in the plasmid-borne mini CRISPR-array. The genome of one randomly chosen colony was sequenced, and indeed a duplication of a fragment of the *hfq* protospacer that should prevent recognition by the CRISPR effector was observed (Supplementary Table 1).

The results presented in Fig. 2C suggest that self-targeting has a bactericidal effect. Previously, we used a similar self-targeting system to study the details of CRISPR action in *E. coli* (*34*). We found that extended regions of DNA flanking the target protospacer were removed due to the Cas3 nuclease/helicase action. We were interested in determining the fate of DNA at and around the targeted protospacer in *C. difficile*. Accordingly, we prepared genomic DNA from ATc-induced cultures 1 hour post-induction, when the drop in viable cell counts was evident (Fig. 2E), and 3 hours post-induction, when growth inhibition of cultures carrying the self-targeting plasmid started to appear (Fig. 2E). Genomic DNA was prepared from each culture and subjected to whole genome sequencing (WGS). The resulting reads were mapped onto the *C. difficile* 630Δ*erm* genome. The overall genome coverage for each culture was between 200 and 300. In the 3 hour induced culture of cells harboring the self-targeting plasmid, a deep drop in the coverage centered at the targeted protospacer in the *hfq* gene was observed. The coverage gradually and symmetrically increased to the mean level of ca. 100 kbp upstream and downstream of the targeted protospacer. The results are very much in line with the *E. coli* data, where self-targeting by a type I-E system was studied (34). Importantly, no decrease in genome coverage in induced cultures of cells harboring the self-targeting plasmid containing the *acrIB2* gene was observed, confirming once again that AcrIB2 is able to abrogate *C. difficile* CRISPR interference. At 1 hour post-induction samples, the decrease in coverage at and around the *hfq* protospacer was minor. Since colony formation by cells collected at this time point is severely decreased (Fig. 2C), we surmise that events leading to the destruction of host DNA have not yet been initiated. Presumably, at the 1-hour time-point, the self-targeting crRNA is not yet produced in sufficient amounts (or did not enter the Cascade complex). However, once such cells are deposited on the surface of the ATc-free medium, sufficient amounts of self-targeting Cascade accumulate and prevent cell division.

In *E. coli,* self-targeting by CRISPR-Cas leads to an SOS response that results in cell filamentation (34). In *C. difficile,* DNA damage also leads to filamentous cell morphology (35, 36). We decided to check the cell morphology of induced self-targeting *C. difficile*. Compared to controls, elongation of *C. difficile* cells carrying the self-targeting plasmid was observed in cultures collected 3 hours post-induction (Supplementary Fig. 2).

We also tested the ability of AcrIB2 to overcome CRISPR interference in a biologically relevant context. φCD38-2 is a prophage of *C. difficile* CD125 strain. We conjugated CD125 and the isogenic R20291 strain that lacks the prophage with a plasmid containing the self-targeting mini-array or an empty vector. Transconjugants were selected in the absence of ATc. Next, transconjugant cultures were serially diluted and spotted on plates with and without the ATc inducer. As can be seen from Fig. 3, the number of viable cells decreased in cultures of R20291 carrying the self-targeting plasmid by at least 10-fold. No such effect was observed in cells that carried the prophage. As expected, no decrease in viable cell counts was observed in ATc-induced R20291 carrying the self-targeting plasmid that also expressed AcrIB2.

**Figure 3.**
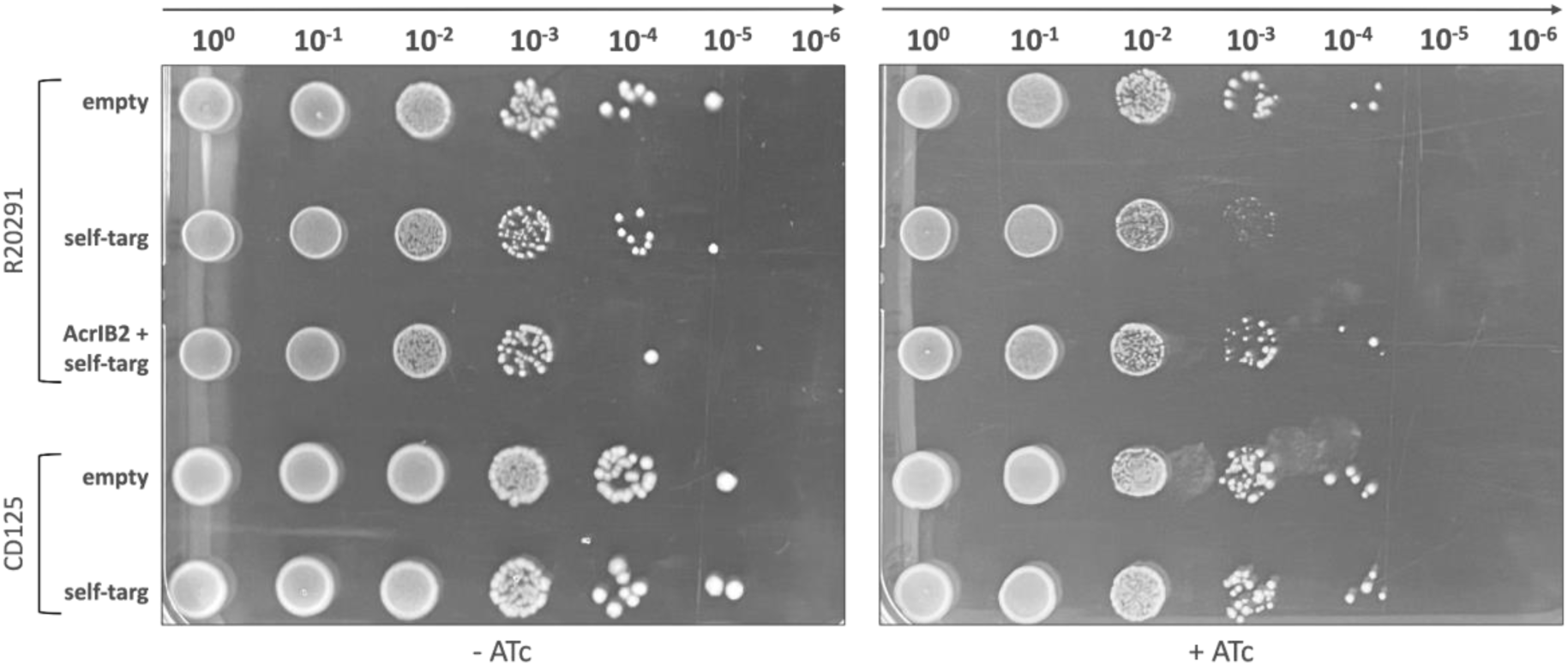
AcrIB2 expressed from a prophage decreases CRISPR interference. Tenfold serial dilutions of transconjugant mixtures of control (“empty”), self-targeting (“self-targ”), or AcrIB2+self-targeting (“AcrIB2+self-targ”) plasmid for R20291 control *C. difficile* strain or CD125 derivative carrying the φCD38-2 prophage were deposited on the BHI agar plates supplemented with Tm in the presence or in the absence of the ATc inducer. A representative result from at least three independent experiments is shown.

### Only one of the two *C. difficile* type I-B *cas* operons is interference-proficient and is targeted by AcrIB2

Most *C. difficile* strains contain at least two *cas* operons per genome (Supplementary Fig. 3). For example, the *C. difficile* 630Δ*erm* strain used in this work carries two (27). The first *cas* operon, *CD2982-CD2975,* is referred to as a “full” *cas* operon and encodes a complete set of proteins necessary for both interference and adaptation. In contrast, the second *cas* operon, *CD2455-CD2451*, referred to as “partial”, lacks genes coding for adaptation proteins Cas1, Cas2, and Cas4 (Supplementary Fig. 3A). Notably, the full *cas* operon is absent in ∼10% of analyzed *C. difficile* strains. The relative contribution of each of the two *cas* operons to CRISPR interference remains unclear.

To determine the contribution of individual *C. difficile* 630Δ*erm cas* operons and identify which one of them is targeted by AcrIB2, we generated mutants lacking either the full *cas* operon (Δfull), the partial one (Δpartial) or both (Δdouble). The strains were conjugated with plasmids carrying the self-targeting mini CRISPR arrays with or without *acrIB2*. Wild-type *C. difficile* 630Δ*erm* was used as a control. Transconjugants were selected on plates without the ATc inducer, and the number of viable cells was determined by comparing cell counts on media with and without the inducer.

All strains formed the same amounts of CFUs in the absence of the inducer, though colonies formed by wild-type and Δfull cells carrying the self-targeting plasmids appeared to be slower (Fig. 4 middle panel), indicting slower growth possibly due to partial self-interference in the absence of the inducer. CRISPR interference in the Δfull mutant was as efficient as in the wild-type control (as judged by the drop of viable cells upon induction of self-interference, Fig. 4 right panel). In contrast, the viability of cells in either the Δpartial or the Δdouble mutant cultures was not affected by induction. Thus, the full *cas* operon is not capable of interference, at least with the self-targeting crRNA used. Expression of *acrIB2* restored the viability of cells in induced self-targeting wild-type and Δfull cultures (Fig. 4, right panel), indicating that the products of the partial operon are inhibited by AcrIB2.

**Figure 4.**
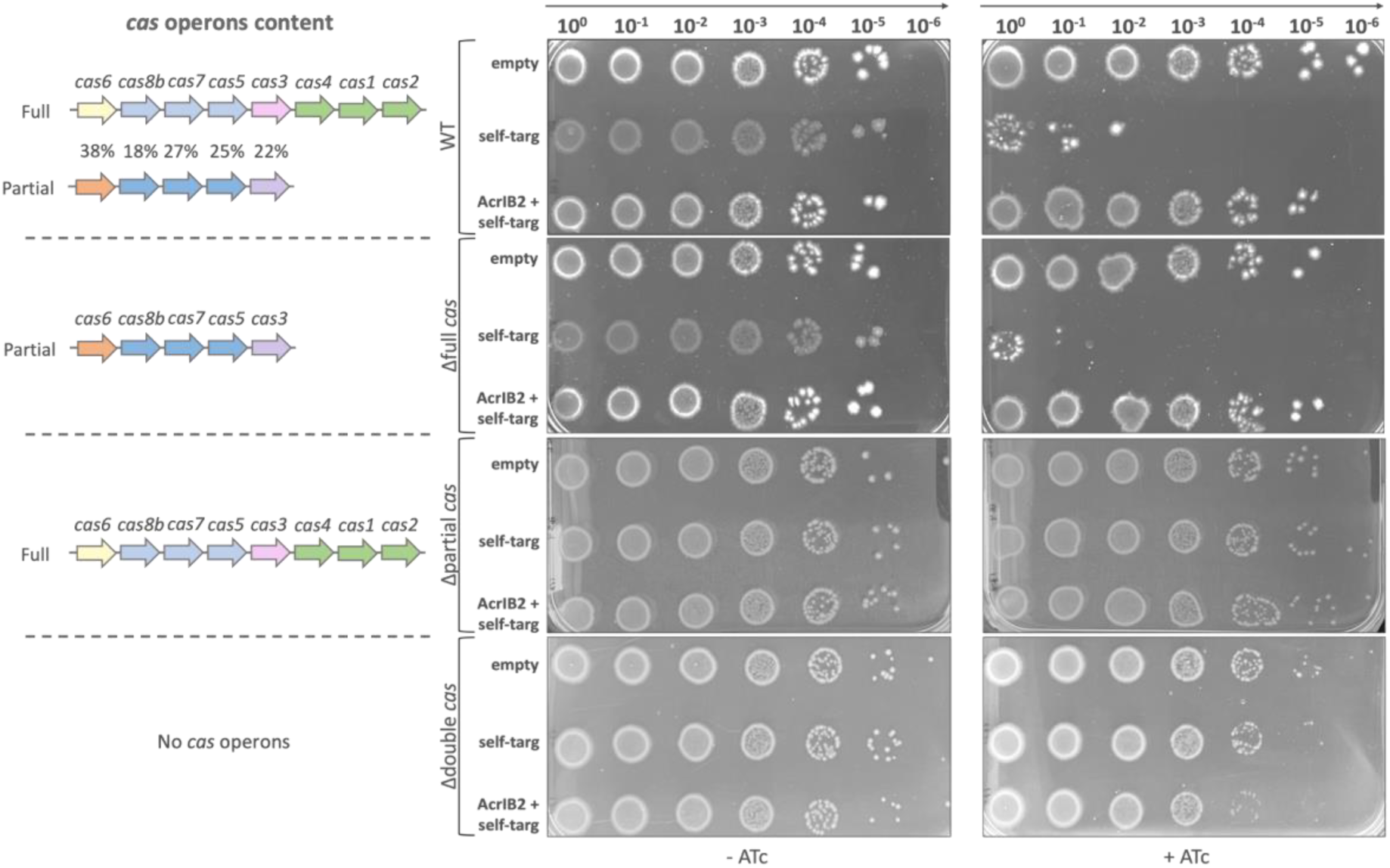
The partial *C. difficile cas* operon is responsible for CRISPR interference and is targeted by AcrIB2. On the left side, *cas* operons content is depicted for each strain. The percentage of amino acid sequence identity of corresponding products between two *cas* operons of *C. difficile 630Δerm* WT are indicated. Middle and right panels show growth of tenfold serial dilutions of indicated cells conjugated with control, self-targeting, and AcrIB2+self-targeting plasmids on the surface of BHI agar plates with or without the ATc inducer.

The finding that the full *cas* operon is apparently non-functional is an unexpected one since sequence analysis of the products of the full operon reveals no potentially inactivating mutations in any of the genes. In the prior study, RNA-seq analysis of *C. difficile 630Δerm* showed that the steady-state levels of transcripts of both *cas* operons are comparable and low in relation to an overall average transcription level under standard laboratory growth conditions with rather uniform coverage detected by both RNA-seq and qRT-PCR (27). To estimate the relative amounts of protein products of both operons, lysates of *C. difficile 630Δerm* were analyzed by liquid chromatography coupled to tandem mass spectrometry (LC-MS/MS). In agreement with the RNA-seq data, the relative quantitative values (total spectrum counts) for Cas proteins were low (between 1 and 21, Supplementary Table 2). For comparison, the relative quantitative values for the most abundant *C. difficile* protein SlpA is more than 3,000, and 150 for RpoA, a subunit of RNA polymerase. Relative quantitative values for subunits of Cascade encoded by the full *cas* operon were consistently 2-3 times lower than for the counterpart encoded by the partial operon. As expected from Cascade stoichiometry, total spectrum counts for Cas7 proteins encoded by each operon were the highest (Supplementary Table 2), adding confidence to our measurements. Perhaps most significantly, the relative quantitative values for Cas3, a protein strictly required for interference, were 20 times higher for the product of the partial operon and minimal (a total spectrum count of 1) for the product of the full operon. We, therefore, speculate that the inactivity of the full operon is due to the low levels of its protein products, most probably, Cas3.

## Discussion

Anti-CRISPR proteins have evolved in response to the co-evolutionary arms race between prokaryotes and their viruses. These proteins exhibit a wide range of structural and functional diversity, and only a small fraction of them have been identified and functionally characterized to date (37). The discovery of Acr proteins has a wide range of applications, including phage therapy of pathogenic bacteria, where Acrs can inhibit the CRISPR-Cas system of the host, thus increasing the ability of the phage to clear the infection (14).

The *C. difficile* CRISPR-Cas system is highly effective in combating MGEs and presumably contributes to the ability of this dangerous pathogen to survive in the phage-rich microbiome of the colon. Indeed, multiple spacers matching the genomes of phages infecting *C. difficile* have been identified (27). All currently identified phages of *C. difficile* are temperate and are capable of either inserting their genetic material into the bacterial genome or exist as episomes (23). Therefore, it is likely that *C. difficile* phages evolved anti-CRISPR mechanisms to protect themselves from CRISPR targeting while in the lysogenic state. However, no such mechanisms have been defined.

In this work, we describe the first anti-CRISPR protein that inhibits the type I-B CRISPR-Cas system of *C. difficile*. The putative anti-CRISPR locus was identified in *C. difficile* phage genomes in the course of a bioinformatic search that used a previously validated AcrIC5 from *Pseudomonas* as bait. We identified a gene coding for a homologous protein next to a gene coding for a potential DNA-binding AP2 domain protein known to be associated with some *acr* loci (20). Using the AP2 domain protein as another bait, putative Acr proteins in several *C. difficile* phages were predicted (Fig. 1B). The putative clostridial Acrs are similar to each other but share no identifiable sequence similarity to known Acrs.

The validated *acr* gene of *C. difficile* phage φCD38-2 (*acrIB2*), together with two unknown-function genes upstream, is located immediately downstream of a long cluster of capsid, DNA packaging, tail, and lysis proteins genes and is transcribed in the same direction (38). Immediately downstream of *acrIB2* is a putative lysogenic conversion region that is transcribed in the opposite direction. The *acrIB2* gene and other upstream genes shown in Fig. 1B are highly transcribed in a stable lysogen carrying the φCD38-2 episome (38). Phages that encode *acrIB2* homologs belong to different morphological classes (sipho- and myoviridae) and likely rely on different developmental strategies. While some phages encode an AP2 domain protein used for the search, others, including the φCD38-2 that encodes the validated AcrIB2 protein, do not (Fig. 1B). Some of the unknown-function genes that are adjacent to *acrIB2* gene homologs in these phages may encode novel Aca proteins. Interestingly, the majority of phages possess a highly conserved gene of an unknown function downstream from *acrIB2* homologous genes. Of particular interest is phage φCD211 (39). Its genome is much larger than the genomes of other phages encoding AcrIB2 homologs. In the immediate neighborhood of its *acrIB2*-like gene, there is an open reading frame coding for a short C-terminal fragment of a Cas3-like protein and a 4-spacer CRISPR array targeting some known *C. difficile* phages (39). It is possible that this locus is used in inter-phage warfare as other prophage-located and prophage-targeting CRISPR arrays in several *C. difficile* strains (27).

Our top hits for the AP2 domain protein encoding gene in *C. difficile* phages are neighbored by genes encoding split AcrIB2 homologs. Presumably, these phages encode either a unique split anti-CRISPR protein or produce a fusion protein as a result of +1 translational frameshifting between gp28 and gp29 ORFs as previously described in other bacteriophages (40–42).

AcrIB2 has a very strong effect on CRISPR interference against conjugating plasmids in the self-targeting model when expressed from an inducible promoter. In a biologically more relevant context of a φCD38-2 lysogenic strain, its effects are milder, increasing survival in a self-targeting model ca. 10-fold. Although this has not been tested in this work, we assume that the protective effects of AcrIB2 in the context of phage infection will also be partial. We attempted to delete the *acrIB2* gene from the φCD38-2 genome. Regrettably, this proved impossible, perhaps because in the φCD38-2 lysogens multiple copies of phage episome exist, making it difficult to select desired clones.

The AcrIB2 protein and its homologs from other *C. difficile* phages have a high number of negatively charged amino acids and aromatic amino acids. Thus, they may act as a DNA mimic (Fig. 5A). The secondary structure prediction indicates a prevalence of alpha-helix motifs in the protein structure (Fig. 5B). The AcrIB2 structure predicted with the AlphaFold tool reveals clustering of negatively charged residues along the long axis of the protein (Fig. 5C), consistent with the DNA mimicry hypothesis regarding the mechanism of action of AcrIB2. The negatively charged positions are conserved among AcriB2 homologs, suggesting their essentiality (Fig. 5A, 5D). In the predicted structure, the position of the split that occurs in cases when an AcrB2 homolog is encoded by two separate genes is located in an unstructured linker (Fig. 5D) and should not prevent the C-terminal fragment of the protein from making tight interactions with the N-terminal part that makes a structurally compact core from which a linker with conserved negatively charged residues (D92, E94, E95, Fig. 5D) protrudes.

**Figure 5.**
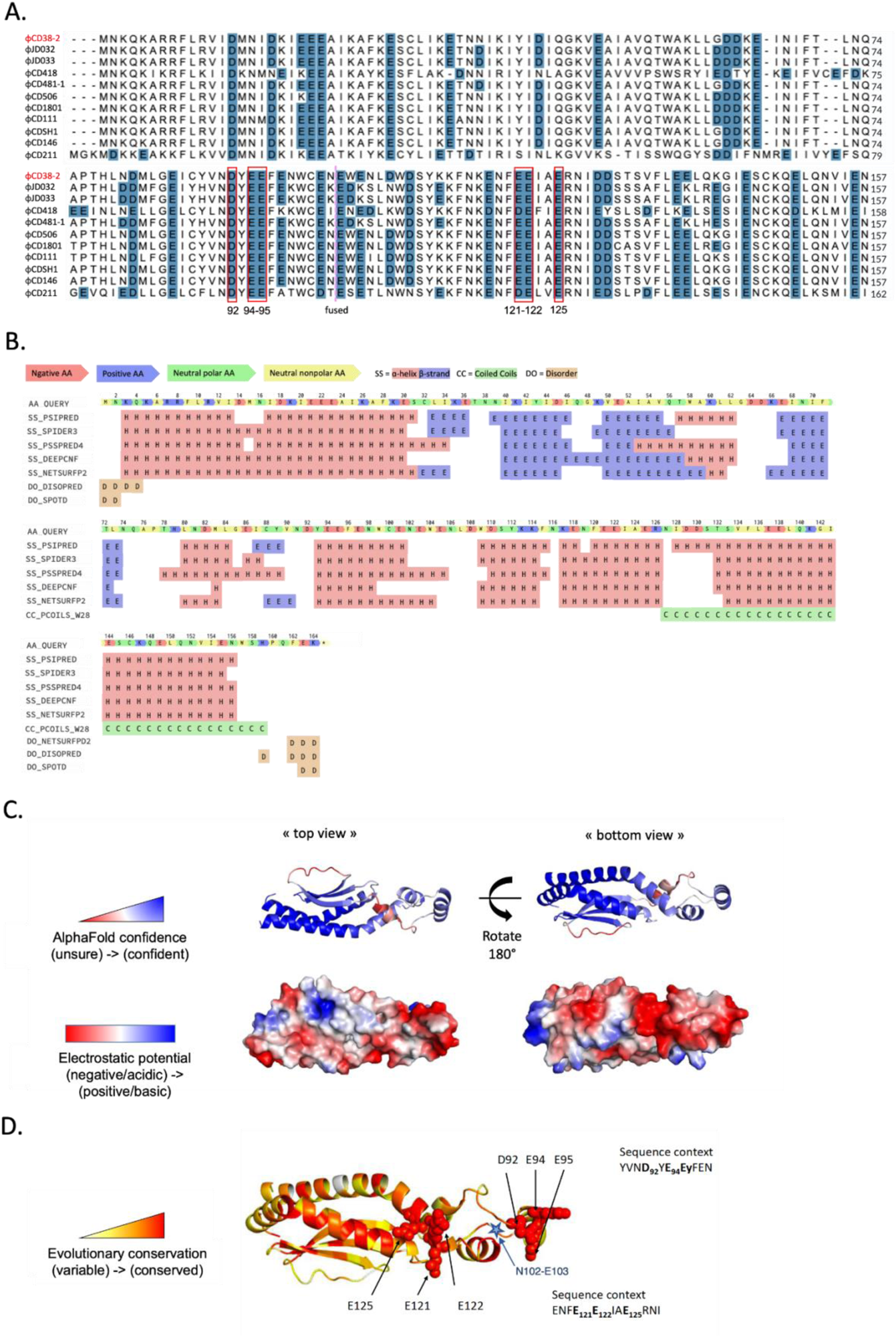
AcrIB2 structure prediction. (A) Alignment of several AcrIB2 homologs performed with UniProt Align tool (43). Negatively charged amino acids are highlighted in blue. Conserved amino acids are marked in red frames. The purple dashed line indicates the location of the split in AcrIB2 homologs encoded by two separate genes. (B) AcrIB2 secondary structure prediction with Quick2D tool (44). (C) The AlphaFold AcrIB2 structure prediction with indicated confidence (as measured by the pLDDT score, from red for low model confidence to blue for high confidence) and electrostatic potential mapped on the surface. (D) Mapping of evolutionary conservation on the AcrIB2 AlphaFold structural model, from white (variable) to red (conserved). Side chains of clustered conserved amino acids are shown in spacefill representation. The blue star indicates the position of a split that occurs in AcrIB2 homologs encoded by two separate genes.

Most *C. difficile* strains contain two *cas* operons, and their individual contribution to interference was not explored before the present study. Surprisingly, our results demonstrate that the mutant lacking the partial *cas* operon exhibited a complete loss of CRISPR interference activity, which indicates that it plays the primary role in CRISPR defense that is inhibited by AcrIB2. When expressed in *E. coli* heterologous host, the full *cas* operon led to a decrease in the transformation rate of CRISPR-targeted plasmid even though the efficiency was rather low as compared to CRISPR interference in natural settings in *C. difficile* (27). Since the partial operon lacks the adaptation module, spacer acquisition must be driven by the products of the full operon. Indeed, we have recently shown that the adaptation module is functional in naive adaptation when expressed from a plasmid (28). Interestingly, both *cas* operons are associated with general stress response SigB-dependent promoters, but we observed a stronger effect of *sigB* mutation on the full *cas* operon expression as compared to partial *cas* operon (45). This differential expression could suggest a potential role of full *cas* operon under stressful conditions. While the function of the interference module of the full *C. difficile cas* operon needs to be specified, it is attractive to speculate that it may be involved in regulatory function in concert with specific crRNAs or, together with the products of the adaptation module, be responsible for primed adaptation.

In conclusion, the identification of a new anti-CRISPR protein targeting *C. difficile* type I-B CRISPR-Cas contributes to a better knowledge of the phage-host relationship and coevolution of defense and counter-defense systems for this important human pathogen and opens interesting perspectives for further developments of applications in biotechnology and health. Apart from its potential applications in phage therapy and phage selection (46), AcrIB2 can also be leveraged as a control for CRISPR-Cas endogenous editing tool (33). Moreover, AcrIB2 holds promise for enhancing the efficacy of the newly developed phage-delivered CRISPR-Cas3 antimicrobial, which triggers the self-elimination of *C. difficile* caused by the activity of the endogenous CRISPR-Cas system(47).

## Material and Methods

### Bioinformatic search of putative anti-CRISPR

The guilt-by-association bioinformatic method was used to identify the putative anti-CRISPR I-B type protein. The method is based on a chain search of homologs of *acr* and *aca* genes using BLAST (48). Uncharacterized ORFs were identified with ORFfinder NCBI (49). The identification of other putative *acr* and *aca* loci in *C. difficile* phages and prophages was made by BLAST search (48). The list of clostridial phages and identified putative Acrs can be found in Supplementary Table 3.

### Plasmid construction

The nucleic acid and amino acid sequences of anti-CRISPRs used in this study are listed in Supplementary Table 4. The list of plasmids used for this study is summarized in Supplementary Table 5. The putative *acr* gene from φCDHM13 phage was cloned into the protospacer, and self-targeting plasmids (pRPF185 derivatives) accompanied by regulatory elements (P_tet_ promoter, RBS, and terminator) in the form of gBlock (dsDNA) from IDT (France). The cloning was achieved through Gibson Assembly by using NEB Gibson Assembly® Master Mix – Assembly (E2611) (50). The resulting constructions were transformed into *E. coli* NEB beta cells (New England BioLabs) and verified by Sanger sequencing. To construct editing plasmids, approximately 800 bp long flanking regions of partial and full *cas* operon of the *C. difficile* 630Δ*erm* strain were amplified by PCR and introduced into the pMSR vector (51) using Gibson assembly reaction (51). The resulting constructions were transformed into *E. coli* NEB beta cells (New England BioLabs) and verified by Sanger sequencing. The list of primers used for this study is summarized in Supplementary Table 6.

### Bacterial strains and growth conditions

All bacterial strains used in this study are listed in Supplementary Table 4. *C. difficile* was cultivated in the anaerobic chamber (Jacomex, France), filled with an atmosphere of 5% H2, 5% CO2, and 90% N2. Both liquid cultures and plate growth were conducted using Brain Heart Infusion (BHI) medium (Difco) at 37°C. When working with strains carrying plasmids, thiamphenicol (Tm) at the final concentration of 15 µg/ml was added to overnight cultures, and 7.5 µg/ml was used for the day cultures. In order to induce the inducible Ptet promoter of pRPF185 derivatives in *C. difficile*, the non-antibiotic analog anhydrotetracycline (ATc) was added at the final concentration of 100 ng/ml. The *E. coli* strains were cultured in LB medium at 37°C supplemented with 100 μg/ml ampicillin (Amp) and 15 µg/ml chloramphenicol (Cm) when required.

### Plasmid conjugation and estimation of conjugation efficiency

All plasmids were transformed into the *E. coli* strain HB101 (RP4). Transformants were further mated with *C. difficile* cells on BHI agar plates for 8h (for *C. difficile* 630) or 24h (for *C. difficile* R20291) at 37°C. Further, *C. difficile* transconjugants were selected on BHI agar plates containing Tm (15 µg/ml), D-cycloserine (Cs) (25 µg/ml), and cefoxitin (Cfx) (8 µg/ml). To estimate conjugation efficiency, after the mating step, *C. difficile* conjugation mixture was serially diluted and plated on BHI agar supplemented with Tm, Cs, and Cfx, or Cs and Cfx only. Then the ratio of *C. difficile* transconjugants to the total number of CFU/ml was estimated.

### Growth assays

*C. difficile* carrying either plasmid maintained in 7.5 µg/ml Tm was grown to an OD_600_ equal to 0.4-0.5, after which ATc inducer was added to a final concentration of 100 ng/ml. Then cultures were either transferred to a 96-well plate to obtain growth curves by using the CLARIOStar Plus machine or serially diluted and plated on BHI + Tm (15 µg/ml) plates at a certain time point and grown overnight before CFU counting. For the drop tests, *C. difficile* carrying either plasmid was serially diluted from starting OD_600_ of 0.4 and spotted on BHI Tm plates (15 µg/ml) with or without ATc inducer (100 ng/ml). Plates were incubated at 37°C for 24h or 48h and photographed.

### Microscopy

For phase-contrast microscopy, *C. difficile* carrying either plasmid maintained in 7.5 µg/ml Tm was grown to an OD_600_ equal to 0.4-0.5, after which ATc inducer was added to a final concentration of 100 ng/ml. After 3 h of incubation at 37°C, 1 ml of culture was centrifuged at 3500 rpm for 5 min, and the pellet was resuspended in 20 µl of sterile H_2_O. Cells were fixed with 1.2% agarose on the slide. Images were captured on a Leica DM1000 microscope using a FLEXACAM C1 12 MP camera with the LAS X software.

### High-throughput sequencing of total genomic DNA

Total genomic DNA was purified by NucleoSpin Microbial DNA Mini kit (Machery-Nagel). For library preparation, the NEBNext® Ultra™ II DNA Library Prep Kit for Illumina (NEB) was used, and the sequencing was carried out on an Illumina platform (NovaSeq 6000). To ensure accurate data analysis, raw reads were trimmed using Trimmomatic v0.39 (NexteraPE-PE.fa:2:30:10 LEADING:3 TRAILING:3 SLIDINGWINDOW:4:15 MINLEN:20). Reads were then aligned to the reference genome using Bowtie2 aligner with end-to-end alignment mode and one allowed mismatch (52). Only reads with unique alignment were retained for further analysis. BAM files were analyzed using the Rsamtools package, and reads with MAPQ scores equal to 42 were selected for downstream coverage analysis and calculating the mean coverage across the genome (34, 53).

### Deletion of *cas* operons in *C. difficile*

An allele-coupled exchange mutagenesis approach described previously (51) was used to delete the partial and full *cas* operons from the *C. difficile* 630Δ*erm* strain. Editing plasmids were conjugated into *C. difficile.* Transconjugants were selected on BHI supplemented with Cs, Cfx, and Tm and then restreaked onto fresh BHI plates containing Tm twice in a row to ensure the purity of the single crossover integrant. The purified colonies were then streaked onto BHI plates with ATc (100 ng/ml) plates to ensure the selection of cells where the plasmid had been excised and lost. If the plasmid was still present, the toxin was produced at lethal levels, and colonies did not form in the presence of ATc. Growing colonies were tested for the success of the deletion by PCR and Sanger sequencing.

### AlphaFold structure prediction

The AcrIB2 amino-acid sequence was used as input of the MMseqs2 homology search program (54) with three iterations against the Uniref30_2202 database to generate a multiple sequence alignment (MSA). This MSA was filtered with HHfilter using parameters (‘id’=100, ‘qid’=25, ‘cov’=50), resulting in 68 homologous sequences, then full-length sequences were retrieved and realigned with MAFFT (55) using the default FFT-NS-2 protocol. Then 5 independent runs of the AlphaFold2 (56) algorithm with 6 recycles were performed with this input MSA and without template search, using a local instance of the ColabFold (57) interface on a local cluster equipped with an NVIDIA Ampere A100 80Go GPU card. Each run generated 5 structural models. The best model out of 25 was picked using the pLDDT confidence score as a metric and used for further structural analysis (pLDDT for this model: 83.5). The qualitative electrostatic surface was generated using PyMOL (58) (local protein contact potential). The evolutionary conservation scores were generated using the AlphaFold2 MSA as an input to the Rate4Site (59) program, which computes the relative evolutionary rate for each site.

## Acknowledgments

We thank Louis-Charles Fortier for stimulating discussions and providing us with the CD125 strain and Ana Margarida Oliveira Paiva for helpful discussions and assistance with microscopy analysis. We also thank Sofia Medvedeva, Anna Shiriaeva for helpful discussions and Sergei Borukhov for advice on protein co-purification. We are grateful to Christophe Le Clainche’s group for providing us with Streptactin reagents. This work has benefited from the facilities and expertise of the Proteomic-Gif (SICaPS) platform of I2BC (Institute for Integrative Biology of the Cell, CEA, CNRS, Université Paris-Saclay, 91198, Gif-sur-Yvette Cedex, France). We thank members of the SICaPS platform for assistance with mass spectrometry analysis. This work was supported by the Institut Universitaire de France (to O. S.), the Institute for Integrative Biology of the Cell, the University Paris-Saclay, Graduate School Life Sciences and Health, and OI MICROBES funding and Vernadski fellowship (to P. M.). This work was also supported by NIH grant RO1 10407 to (K. S.), the Russian Science Foundation grant 19-14-00323, and the Ministry of Science and Higher Education of the Russian Science Federation agreement no. 075-10-2021-114.

## Notes

### Competing Interest Statement

The authors have declared no competing interest.

