## Supplementary material for "Identification of an anti-CRISPR protein that inhibits the CRISPR-Cas type I-B system in *Clostridioides difficile*"

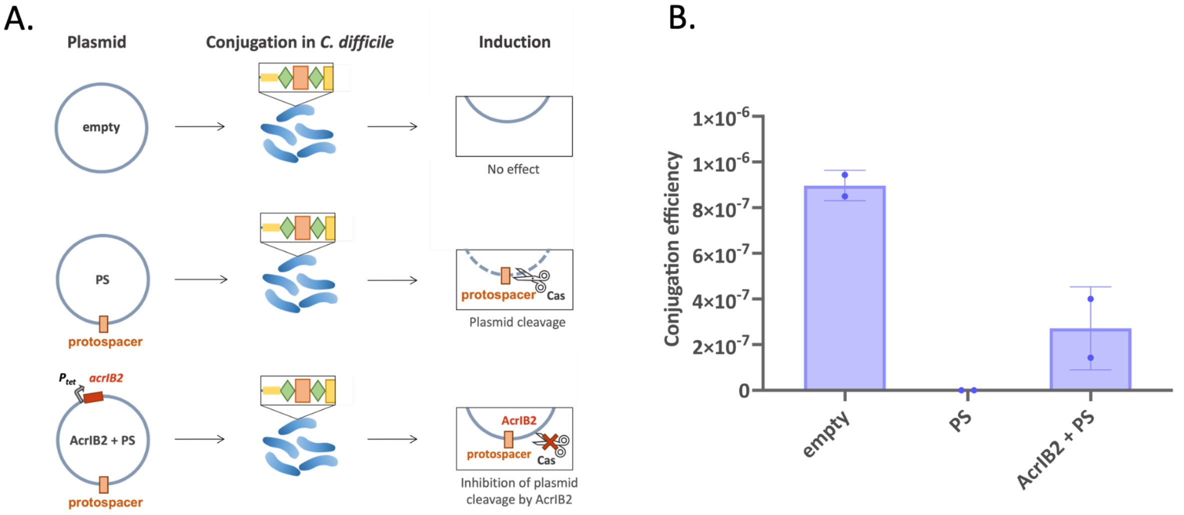


**Supplementary Figure 1. Anti-CRISPR protein AcrIB2 partially inhibits CRISPR interference of pRPF185Δgus derivative plasmid carrying the protospacer.**

(A) A plasmid loss strategy to reveal anti-CRISPR activity of plasmid-borne genes relies on a plasmid that carries a protospacer sequence (orange box) targeted by the crRNA spacer of the *C. difficile* 630Δerm CRISPR3 array. Green rhombi indicate CRISPR repeats, the blue rectangle indicates the spacer, the leader sequence is indicated in yellow. “PS” stands for protospacer plasmid, and “AcrIB2+PS” stands for protospacer plasmid with ATc-inducible *acrIB2* gene (red box). The control plasmid is referred to as “empty” vector.

(B) The conjugation efficiency of plasmids used for plasmid loss strategy is shown. Experiments were repeated at least two times, and error bars represent the standard deviations between two replicates.

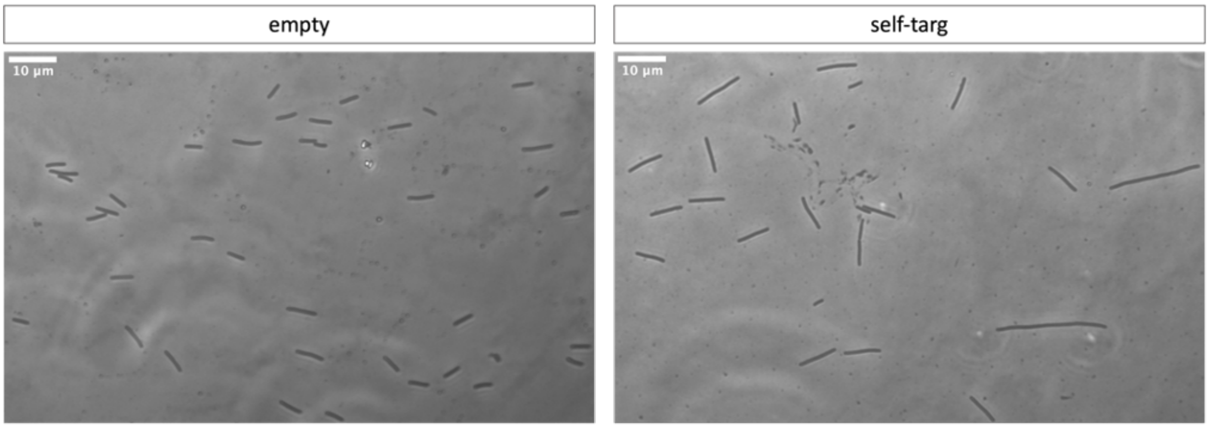


**Supplementary Figure 2.** The micrograph on the right shows a representative field of cells (x40 magnification) illustrating the effect of 3-hour induction of self-targeting plasmid-borne CRISPR array transcription on the cell morphology. The micrograph on the left shows cells carrying the control plasmid with no mini CRISPR array grown in similar conditions.

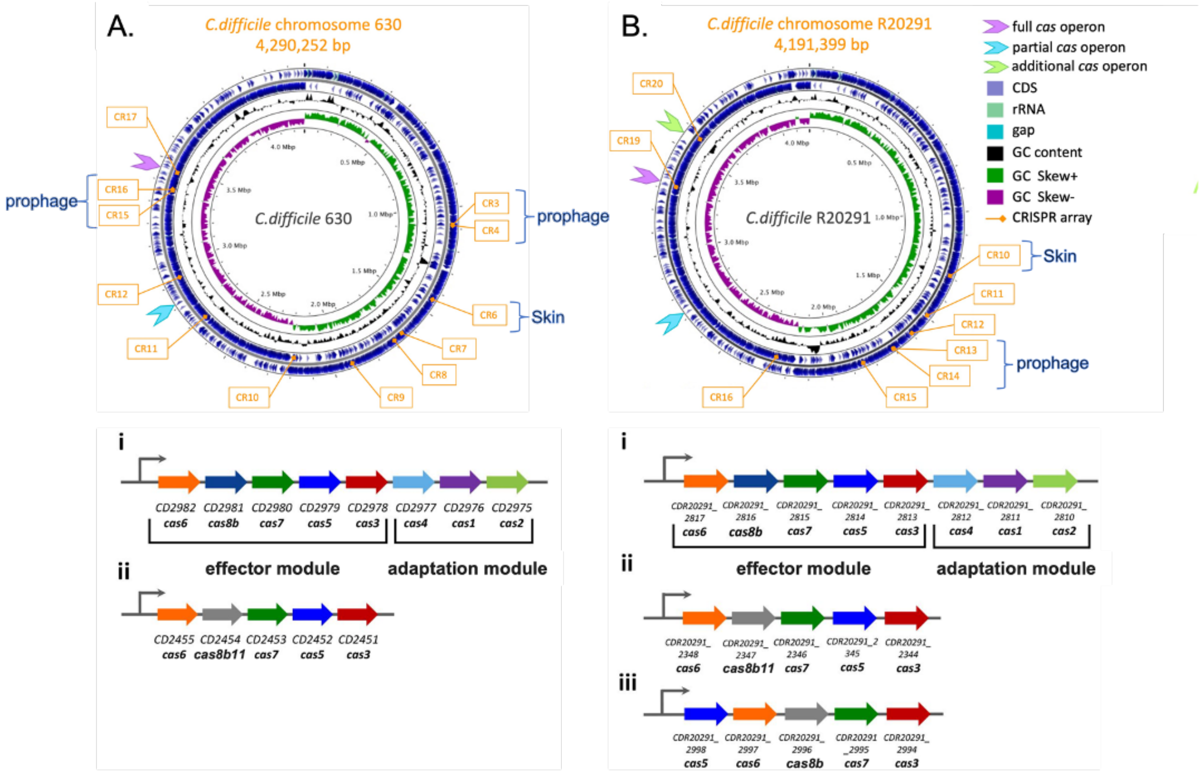


**Supplementary Figure 3.** Schematic view of CRISPR-Cas organization in the chromosome of *C. difficile* strains 630 (A) and R20291 (B). CRISPR arrays (CR) are numbered according to the CRISPRdb database (60). The locations of associated *cas* operons and prophage regions are indicated. The organization of the *cas* operons in strain 630 (A) and R20291 (B) are indicated with roman numerals, where i – full operons; ii – partial operons; iii – an additional operon. Functional modules are marked with brackets. The same color was used for homologous *cas* genes. (27)

**Supplementary Table 1.** Mutations found in genomic DNA of escapers colonies of *C. difficile* 630Δ*erm* WT and Δfull *cas* operon mutant.

| **Strain** | **Mutation** | **Position** | **Gene/Region** | **Translation** |
|---|---|---|---|---|
| WT | Insertion of A | 255688 | CD630_01960 (Nickel carrier?) | Frameshift |
| | A>C | 694997 | Intergenic region | - |
| | G>C | 1125053 | Spacer1 of CRISPR3 | - |
| | A>C | 2069420 | CD630_17860 | N290H |
| | C>T | 2663884 | Leader seq of CRISPR11 | - |

| | | | | |
|---|---|---|---|---|
| | C>G | 3397588 | Spacer1 of CRISPR16 | - |
| | T>G | 4034625 | CD630_34420 | F63V |
| | Deletion (AAAAGTGCTAAAGCACGGCGCCAACGGATAAATCCCGTTGCTT) | 4150289 | Intergenic region | - |
| Δfull *cas* operon | T>C | 26911 | Intergenic region | - |
| | G>T | 26912 | Intergenic region | - |
| | Insertion (+TAAATATATAGT) | 2276149 | PPS (*hfq* gene) | Modifies the 3' of the PPS sequence |
| | T>G | 2316896 | CD630_20061 | STOP codon |
| | Deletion (AAAAGTGCTAAAGCACGGCGCCAACGGATAAATCCCGTTGCTT) | 3152458 | Intergenic region | - |
| | T>A | 3397586 | Spacer1 of CRISPR16 | - |
| | A>G | 3682242 | Intergenic region | - |
| | T>C | 3682243 | Intergenic region | - |
| | G>A | 3682337 | Intergenic region | - |
| | T>C | 3682342 | Intergenic region | - |
| | A>C | 3682343 | Intergenic region | - |
| | A>G | 3682346 | Intergenic region | - |
| | T>C | 3682347 | Intergenic region | - |
| | T>C | 3682349 | Intergenic region | - |
| | Insertion of A | 3972603 | Intergenic region | - |

| | | | | |
|---|---|---|---|---|
| | T>A | 3999236 | Intergenic region | - |

**Supplementary Table 2.** The relative quantitative values of protein products of both *cas* operons in *C. difficile* 630Δ*erm* found by liquid chromatography coupled to tandem mass spectrometry (LC-MS/MS).

| Cas protein | Molecular Weight | Number of unique peptides | | | Quantitative values (Total spectrum count) | | | Average |
|---|---|---|---|---|---|---|---|---|
| | | W1 | W2 | W3 | W1 | W2 | W3 | |
| **Partial *cas* operon** | | | | | | | | |
| Cas6 | 29 kDa | 4 | 4 | 6 | 4 | 4 | 7 | **5** |
| Cas8b11 | 58 kDa | 10 | 7 | 10 | 14 | 8 | 11 | **11** |
| Cas7 | 39 kDa | 11 | 12 | 13 | 19 | 21 | 22 | **21** |
| Cas5 | 31 kDa | 6 | 3 | 6 | 6 | 3 | 6 | **5** |
| Cas3 | 94 kDa | 15 | 14 | 14 | 20 | 18 | 20 | **19** |
| **Full *cas* operon** | | | | | | | | |
| Cas6 | 29 kDa | 1 | | 1 | 1 | | 2 | **2** |
| Cas8b | 74 kDa | 5 | 4 | 3 | 7 | 7 | 4 | **6** |
| Cas7 | 35 kDa | 6 | 6 | 7 | 6 | 8 | 8 | **7** |
| Cas5 | 27 kDa | 2 | 3 | 1 | 2 | 3 | 2 | **2** |
| Cas3 | 90 kDa | 1 | 1 | 1 | 1 | 1 | 1 | **1** |

**Supplementary Table 3.** List of clostridial phages and identified putative Acrs.

| Phage | Putative *acr* gene name | Gene characteristics | Functionality | Reference for phage |
|---|---|---|---|---|
| AB012111.1 | | | | (61) |

| | | | | |
|---|---|---|---|---|
| c-st | | | | (62) |
| CDKM15 | | | | (63) |
| CDKM9 | | | | (63) |
| CDMH1 | | | | (64) |
| CDMH11 | *phiCDHM11_gp29&*<br>*phiCDHM11_gp30* | double orf | | (64) |
| CDSH1 | *CDHS1_28* | single orf | | (64) |
| CD1801 | *QVW56634.1* | single orf | | (65) |
| Clo-PEP-1 | | | | (66) |
| CPS1 | | | | (67) |
| CPS2 | | | | (68) |
| CpV1 | | | | (66) |
| D90210.1 | | | | (69) |
| FN668943.1 | | | | (70) |
| JD032 | *JD032_24* | single orf | | (71) |
| JD033 | *JD033_49* | single orf | | (66) |
| phiCD211 | *phiCD211_20148* | single orf | | (39) |
| HM_T | | | | (72) |
| HM2 | | | | (66) |
| phiCDHM13 | *phiCDHM13_gp26 &*<br>*phiCDHM13_gp27,*<br>*phiCDHM13_gp28 &*<br>*phiCDHM13_gp29* | double orf | Non-functional | (64) |
| phi027 | | | | (38) |
| phi24R | | | | (73) |
| phi34O | | | | (74) |
| phi3626 | | | | (75) |
| phi8074-B1 | | | | (76) |
| phi9O | | | | (74) |
| phiC2 | | | | (77) |
| phiC2_isolate_584 | | | | (77) |
| phiC2_isolate_RW11 | | | | (77) |
| phiC2_isolate_RW2 | | | | (77) |

| | | | | |
|---|---|---|---|---|
| phiCD111 | *phiCD111_20028* | single orf | | (27, 78) |
| phiCD119 | | | | (79) |
| phiCD146 | *phiCD146_20027* | single orf | | (27, 78) |
| phiCD24-1 | | | | (27, 78) |
| phiCD24-2 | | | | (27, 78) |
| phiCD27 | | | | (80) |
| phiCD38-2 | *phiCD38-2_gp27 (acrIB1)* | single orf | Functional | (31) |
| phiCD481-1 | *phiCD48101_20029* | single orf | | (27, 78) |
| phiCD505 | | | | (27, 78) |
| phiCD506 | *phiCD506_20031* | single orf | | (27, 78) |
| phiCD52 | | | | (27, 78) |
| phiCD630-1 | | | | (26) |
| phiCD630-2 | | | | (26) |
| phiCD6356 | | | | (81) |
| phiCDHM11 | *phiCDHM11_gp26 & phiCDHM11_gp27, phiCDHM11_gp28 & phiCDHM11_gp29* | double orf | | (82) |
| phiCDHM14 | *phiCDHM14_gp26 & phiCDHM14_gp27, phiCDHM14_gp28 & phiCDHM14_gp29* | double orf | | (83) |
| phiCDHM19 | | | | (83) |
| phiCP13O | | | | (74) |
| phiCP26F | | | | (74) |
| phiCP7R | | | | (84) |
| phiCPV4 | | | | (84) |
| phiCT19406A | | | | (85) |
| phiCT19406B | | | | (85) |
| phiCT19406C | | | | (85) |
| phiCT453A | | | | (85) |
| phiCT453B | | | | (85) |
| phiCT9441A | | | | (85) |
| phiCTC2A | | | | (85) |

| | | | | |
|---|---|---|---|---|
| phiCTC2B | | | | (85) |
| phiCTP1 | | | | (85) |
| phiMMP01 | | | | (27, 86) |
| phiMMP02 | | | | (27, 86) |
| phiMMP03 | | | | (27, 86) |
| phiMMP04 | | | | (27, 86) |
| phiS63 | | | | (66) |
| phiSM101 | | | | (87) |
| phiZP2 | | | | (84) |
| susfortuna | | | | (88) |
| vB_CpeS-CP51 | | | | (89) |

**Supplementary Table 4.** Anti-CRISPR sequences used in this study, gBlocks, and regulatory elements.

| **Name** | **Sequence (5’-3’)** |
|---|---|
| *acrIB2* gene (471 bp) | ATGAATAAACAAAAAGCTAGAAGATTTTTAAGAGTTATAGATATGAATATAGATA<br>AAATAGAGGAAGAAGCTATAAAAGCTTTTAAAGAAAGTTGTTTAATCAAAGAGA<br>CTAATAATATAAAAATTTATATCGATATACAAGGAAAAGTTGAAGCGATAGCAGT<br>TCAAACTTGGGCTAAACTTTTAGGTGATGACAAAGAAATTAATATTTTCACATTAA<br>ATCAAGCGCCAACTCATTTAAACGATATGCTTGGAGAAATTTGTTACGTAAACGA<br>TTATGAAGAATTTGAAAATTGGTGTGAAAATGAGTGGGAAAATTTGGATTGGGAT<br>AGTTATAAAAAATTCAATAAAGAAAATTTCGAAGAAATTGCTGAAAGAAATATAG<br>ACGATAGCACATCAGTTTTTTTAGAAGAATTACAAAAAGGCATTGAAAGTTGTAA<br>ACAAGAATTGCAAAATGTAATTGAAAATTAA |
| AcrIB2 protein (156 aa; 20 kDa) | MNKQKARRFLRVIDMNIDKIEEEAIKAFKESCLIKETNNIKIYIDIQGKVEAIA<br>VQTWAKLLGDDKEINIFTLNQAPTHLNDMLGEICYVNDYEEFENWCENEWE<br>NLDWDSYKKFNKENFEEIAERNIDDSTSVFLEELQKGIESCKQELQNVIEN |
| **gBlocks with artificial regulatory elements upstream and downstream** | |
| φCDHM13_gp26 (659bp; CDS 450bp) | AAGCACTGATTAGTACTATAACTCAATATAAGCATATCCCCTGGACTTCATGAAA<br>AACTAAAAAAAATATTGACACTCTATCATTGATAGAGTATAATTAAAATAATAAA<br>CAAAGGGGGATATAAAAATGGAAAAATTCATCAGACTTGATTACGATAAGGGCTT<br>TAGAGGAAAAGAACATGTAAGTTCTGCAACTGGAGATGGAGAACATTTTGAAGC<br>AGGAATTAGTTGTTATAAAATAAGTAAAGAAAAATGTGTTGATGCTATAATAAAT |

| | |
|---|---|
| | TTATGTGAATATTGGTTTGAATTTGCAGGTGAATGTCAATTCAAAGATTTCGATAT<br>AAATATTTTTGAAGGATGCTATGTAGGTGAGGGGGCTAGTTACGAAGATTTAGCC<br>ACTTGTGAGAAACATTTATATACTGTAGATGGTTCTTTATTCAATGAAGTTTATGA<br>CTTATATTATATGCATGAAACATACTTAGAAGAAAATGGAAATGTTGAAGAATTA<br>GAAGAAAACTACAAAGATGAATATATAACAACAGAAGAATTTGAAACTAAGATA<br>AAAGAAATGTTTATAAAATATCTGTAGTAAATATTTATAGAGGCTATAAATAGCC<br>TCTATTTTATGTGACTGTATCGTAACTAGAGAACCAAACGACGGAAAAAGCGAT |
| φCDHM13_gp27 (846bp; CDS 636bp) | AAGCACTGATTAGTACTATAACTCAATATAAGCATATCCCCTGGACTTCATGAAA<br>AACTAAAAAAAATATTGACACTCTATCATTGATAGAGTATAATTAAAATATAATC<br>AAAAGGAGGATATAAAAAATGAAAATTGGAGATAAATTTGAAAATCTTACAATA<br>CTAGATATAGAACAAAAAAATGGCAGGAAATATTGCTTATGTAAATGTAGTAATT<br>GCAATAATGAAAAATGGATAAGGGCAGATAGTTTAAAAAGAATAAAAGCATGTG<br>GATGCATGAAAAGTAACACACAGTTTAAACAAAATGATTTAACAGGCAAAAAGTT<br>TGGCAGATTAACAGCAATAAAGAACACTAATAAAAAAGCTAAAAGTGGTCACTA<br>TATTTGGGCTTGTAAGTGTGATTGTGGAAATGAGATTAAGACAGCAGAAAACAAT<br>TTAACTACTGGTAGAACTAAATCATGTGGATGCTTGAAAAAGGAATCTAATATAA<br>AAAATGCAAAGATAGCGTTAAAAGTACATAAAGAAAAAAATATTATTGATGATA<br>CAAATCTATCTATTATAAAAAAGACAGAAGCGTATTCTAATTCAAAAACTAAAAT<br>TAGAGGCGTTTCTTGGAATAAAGAAAAAAGAAAATACTGTGCACAAATAGAATTT<br>AAGAAAATGCATTATAATCTAGGATACTATGACAACATAAGAGAAGCAGAAGAA<br>GCATACAAAAAAGCCAAAGAAAAATTCTTAAAAGAAATAAACGGAAAAGTTTAA<br>TAAATATTTATAGAGGCTATAAATAGCCTCTATTTTATGTGACTGTATCGTAACTA<br>GAGAACCAAACGACGGAAAAAGCGAT |
| φCDHM13_gp28 (525bp; CDS 315bp) | AAGCACTGATTAGTACTATAACTCAATATAAGCATATCCCCTGGACTTCATGAAA<br>AACTAAAAAAAATATTGACACTCTATCATTGATAGAGTATAATTAAAATATAATC<br>AAAAGGAGGATATAAAAAATGAATAAACAAAAAGCTAGAAGATTTTTAAGAGTT<br>ATAGATATGAATATAGATAAAATAGAAGAAGAAGCTATAAAAGCTTTTAAAGAA<br>AGTTGTTTAATCAAAGAGACTAATGATATAAAAATTTATATCGATATACAAGGAA<br>AAGTTGAAGCGATAGCAGTTCAAACTTGGGCTAAACTTTTAGATGATGACAAAGA<br>AATTAATATTTTCACATTAAATCAAGCACCAACTCATTTAGACGATATGTTTGGAG<br>AAATTTACCATGTAAATGATTATGAAGAGTTCGAAAAATGGTGTGAAAAAGGAAG<br>ATAATAAATATTTATAGAGGCTATAAATAGCCTCTATTTTATGTGACTGTATCGTA<br>ACTAGAGAACCAAACGACGGAAAAAGCGAT |
| φCDHM13_gp29 (384bp; CDS 174bp) | AAGCACTGATTAGTACTATAACTCAATATAAGCATATCCCCTGGACTTCATGAAA<br>AACTAAAAAAAATATTGACACTCTATCATTGATAGAGTATAATTAAAATATAATC<br>AAAAGGAGGATATAAAAAGTGAAAAAGGAAGATAAAAGTTTGAATTGGGATAGT<br>TATGAAAAATTCAATAAAGAAAATTTCGAAGAAATCGCTGAAAGAAATATAGAT<br>GATAGCTCATCAGCTTTTTTAGAAAAATTACGTGAAGGCATTGAAAATTGTAAAC<br>AAGAATTACAAAATATAGTTGAAAATTAATAAATATTTATAGAGGCTATAAATAG<br>CCTCTATTTTATGTGACTGTATCGTAACTAGAGAACCAAACGACGGAAAAAGCGA<br>T |
| **Regulatory elements in gBlocks** | |

| | |
|---|---|
| Left homology arm | AAGCACTGATTAGTACTATAACTCAATATAAGCATATCCC |
| $P_{tet}$ promoter | CTGGACTTCATGAAAAACTAAAAAAAATATTGACACTCTATCATTGATAGAGTATAATTAAAATA |
| RBS | TAATCAAAAGGAGGATATAAAAA |
| Terminator | TAAATATTTATAGAGGCTATAAATAGCCTCTATTTTATGTGA |
| Right homology arm | CTGTATCGTAACTAGAGAACCAAACGACGGAAAAAGCGAT |

**Supplementary Table 5.** Bacterial strains and plasmids used in this study.

| **Strain** | **Genotype** | **Source** |
|---|---|---|
| ***E. coli*** | | |
| NEB-10 beta | *Δ(ara-leu) 7697 araD139 fhuA ΔlacX74 galK16 galE15 e14- ϕ80dlacZΔM15 recA1 relA1 endA1 nupG rpsL (StrR) rph spoT1 Δ(mrrhsdRMS-mcrBC)* | New England Biolabs |
| HB101 (RP4) | *supE44 aa14 galK2 lacY1 Δ(gpt-proA) 62 rpsL20* (StrR)*xyl-5 mtl-1 recA13 Δ(mcrC-mrr) hsdSB (rBmB-)* RP4 (Tra+ IncP ApR KmR TcR) | Laboratory stock |
| ***C. difficile*** | | |
| 630Δ*erm* | Sequenced reference strain Δ*ermB* | Laboratory stock (90) |
| R20291 | PCR-ribotype 027 epidemic strain | Laboratory stock |
| CD125 | R20291 carrying φCD38-2 prophage | (91) |
| CDIP741 | 630Δ*erm ΔCD2975-2982* (full *cas* operon) | (45) |

| | | |
|---|---|---|
| CNRS_CD270 | 630Δ*erm* Δ*CD630_2455* (partial *cas* operon) | This work |
| CNRS_CD564 | 630Δ*erm* Δ*CD2975-2982* (full *cas* operon) and Δ*CD630_2455* (partial *cas* operon) double mutant | This work |
| **Plasmids** | | |
| pDIA6103 | pRPF185Δ*gus* vector derivative | (26, 32) |
| pDIA6435 | pRPF185Δ*gus* with the 5' CCA-PAM protospacer, corresponding to the spacer1 from 630Δ*erm* CRISPR3 array | (28) |
| pDIA6555 | pDIA6103 carrying the *hfq* gene targeting CRISPR mini-array with the partial leader sequence under the control of P*tet* promoter | (33) |
| p074 | pDIA6555 + *acrIB2* | This work |
| **Editing Plasmids** | | |
| pMSR | pMSR and plasmid for chromosomal manipulation in *Clostridioides difficile* | (51) |
| p028 | pMSR14 Δ*cas* CD630/2451-2455 partial operon | (45) |
| p233 | pMSR14 + Δ*cas* CD630/2975-82 full operon | This work |

**Supplementary Table 6.** List of primers used in this study.

| **Name** | **Sequence** | **Purpose** |
|---|---|---|
| JP403 | CTGTATCGTAACTAGAGAACCAAAC | Linearization of pRPF185-derived plasmids |
| JP404 | GGGATATGCTTATATTGAGTTATAGTAC | |
| AS005 | agtactataactcaatataagcatatcccTTGACTTAGATATTCATATAGATTATAATATAAAAATGGAGGATATAAAAAATGAATAAAC | |

| | | |
|---|---|---|
| AS006 | ccgtcgtttggttctctagttacgatacagCATAATAACTAAACATTAAAAAACAGTGT | Cloning the *acrIB2 (gp27)* into pDIA6103 |
| AS007 | taattaaaatataatcaaaaggaggatataaaaaatgaatAAACAAAAAG | |
| AS008 | ATTCATTTTTTATATCCTCCTTTTGATTA | |
| PM_42 | GTATAAGTCTCCTCTCAATCATC | *hfq* interference screening |
| PM_43 | ATGCAAAACGTCAGATAACATGG | |
| PM_44 | CTTTCCAGGTTGTATAGTTG | |
| PM_45 | GTTCAATTTAATCTTGGGAGGGTAC | |
| PM_69 | cggtgttttttgttaccctaagtttGAGATAGAAATTATTTTTATTATACGTTTTTTTG | *cas* deletion *CD630_2982* full operon |
| PM_70 | gttaatatattaattgagagaggtgatattatgTAGGTTTGCAGTGAGCGATATTTATGC | |
| PM_71 | gcataaatatcgctcactgcaaacctaCATAATATCACCTCTCTCAATTAATATATTAAC | |
| PM_72 | ggtcatgagattatcaaaaaggagtttGATAGTGAAGTTTCAATTCTAGATATGATA | |
